# Apples-to-apples transcriptomics: a parent-of-origin-resolved ‘Granny Smith’ genome reveals how reference choice shapes biological interpretation

**DOI:** 10.64898/2026.11.17.767359

**Authors:** Heidi Hargarten, Huiting Zhang, Arild Arifin, Harpreet Kaur, Iris Lange, Dongyun Lee, Zhaslan Akhmetov, Ananta Devkota, Ekom-Obong Idio, Cameron March, Bayley McDonald, Taylor Mcnees, Bipin Neupane, Oluwadamiola Olanrewaju, Sadikshya Poudel, Endeshaw Simie, Itsuhiro Ko, Stijn Vanderzande, Dave Rudell, Alex Harkess, Stephen Ficklin, Loren Honaas

## Abstract

0.

The accuracy and reliability of RNA-seq–based transcriptome profiling inherently depends on how well the reference genome represents the genotype being studied, particularly in highly heterozygous, clonally propagated crops with many genetically distinct cultivars, such as apple. Here, we present a chromosome-scale, parent-of-origin-resolved genome assembly and highly complete annotation for ‘Granny Smith’ (assembly BUSCO = 98.6% both haplotypes; annotation BUSCO = 98.7-98.9%). To test how reference genome choice influences transcriptomic interpretation, we mapped RNA-seq data from sun- and shade-exposed ‘Granny Smith’ peel independently to three different apple genome references: GDDH13 (earlier-generation), ‘WA 38’ (modern-generation), and both ‘Granny Smith’ haplotypes (modern and cultivar matched). Despite small difference in unique mapping rate, transcriptome analysis using the modern references showed greater agreement in differentially expressed gene families than GDDH13, while the cultivar-matched ‘Granny Smith’ references recovered additional functional responses associated with oxidative stress, cuticle biology, and other processes relevant to sun exposure and cold storage. Importantly, many responses were consistent with previously identified metabolomic patterns from the same experimental material, providing independent biological support for signals recovered with the matched reference. Reference-dependent differences were greatest for the subtle sun-exposure × storage interaction, indicating reference limitations become more consequential as biological contrasts become finer. Orthogroup-based analyses further showed that both reference-generation differences and paralog representation contribute to cross-genome disagreement. Together, these results demonstrate that seemingly modest differences in mapping performance can substantially alter biological conclusions and highlight the value of high-quality, genotype-matched references for resolving cultivar-specific stress and postharvest physiology in heterozygous perennial crops.

**Article Summary:** This study presents a high-quality ‘Granny Smith’ apple genome, with each parent’s copy assembled separately, and tests how reference genome choice affects interpretation of gene activity data. The authors compared sun-exposed and shaded apple peel during storage, using three reference genomes. Though reads mapped nearly equally well to each genome, the biological processes identified in differential expression (DE) and functional enrichment (FE) analyses differed substantially. These differences grew as the DE comparisons became subtler. Genome assembly generation method and how related gene copies were represented contributed to discrepancies in DE/FE outcomes based on reference genome choice.

## 1. Introduction

Genomic resources for highly heterozygous pome fruit, such as apple (*Malus domestica*), are important for understanding cultivar-specific responses to growing conditions and postharvest treatments. Apples generate $3.2 billion in annual U.S. farm-gate revenue plus an estimated $23 billion in downstream economic output ^1^, and a global revenue estimated at $78 billion ^2^. This economic importance rests on substantial cultivar diversity: over 100 genetically distinct cultivars are commercially produced in the United States alone ^3^, with many different and wide ranging characteristics such as color, flavor, texture, storability, disease resistance, and growth habits, to name a few. These genetic differences also shape differences in management practices and postharvest disorders susceptibility ^4^. For example, ‘Gala’ is prone to softening during postharvest storage ^5,6^, ‘Honeycrisp’ to bitterpit ^7^ and chilling injuries ^8,9^, and cultivars such as ‘Fuji’ and ‘Scilate’ to injury under controlled atmosphere storage ^10,11^.

Originating in Australia, ‘Granny Smith’ (**Figure 1A**) is now the fifth most widely produced cultivar globally ^12–14^ and the only widely-grown green-skinned cultivar, with an estimated global annual production at 1.3–1.5 million metric tons ^15–22^. Its commercial importance and cultivar-specific susceptibility to physiological disorders exemplify the need for genotype-specific genomics resources. Among these disorders, those associated with sun and heat exposure are a growing concern as extreme heat events become more frequent ^23^, compounding financial pressure facing the U.S. apple industry ^24,25^.

**Figure 1.**
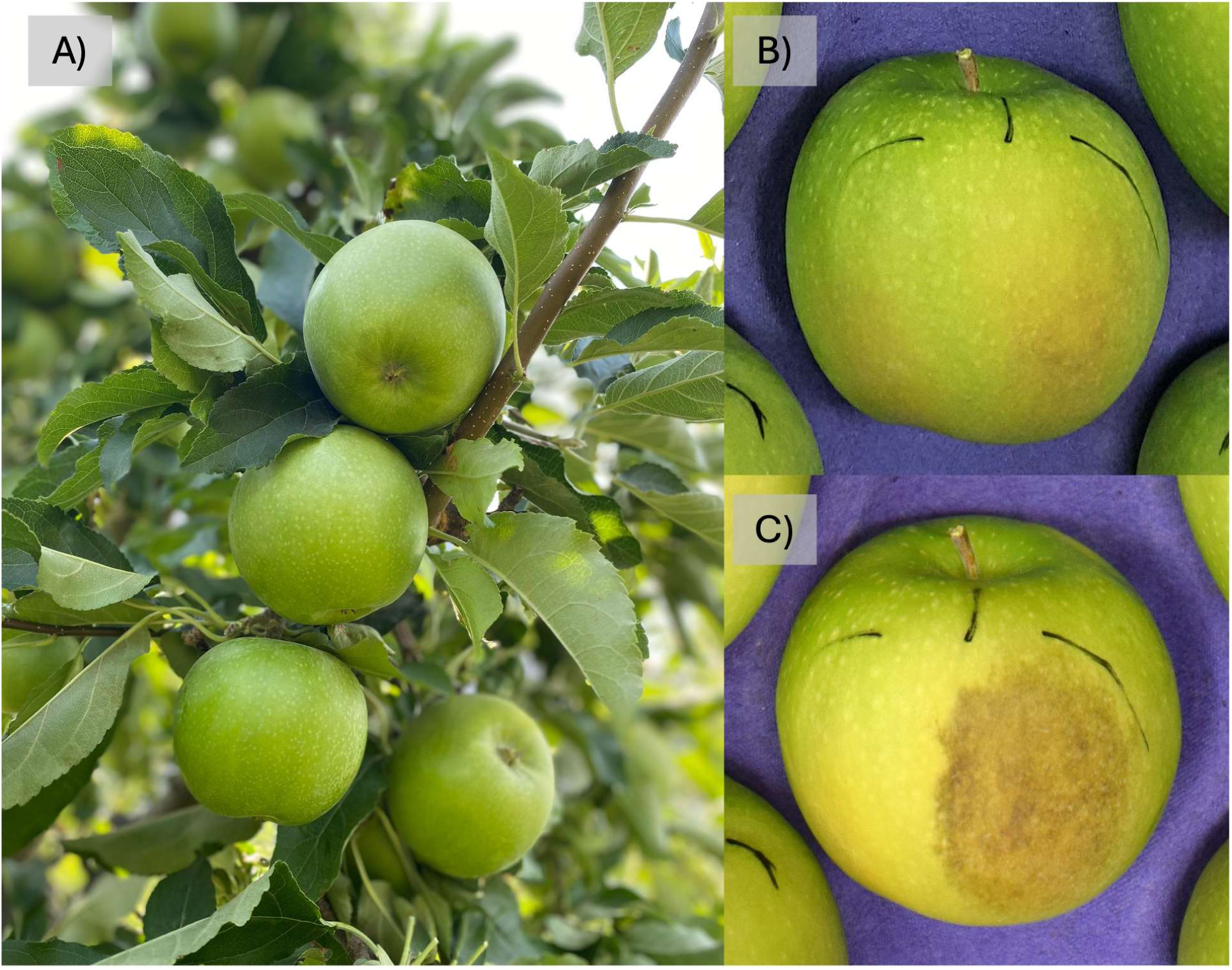
*Malus domestica* ‘Granny Smith’. A) A tart, firm, green apple originating from a chance cross between ‘Cleopatra’ and ‘Stone Pippin’ (C. Peace personal communication) in Australia. ‘Granny Smith’ is a slow maturing cultivar that is susceptible to a variety of postharvest physiological issues triggered by preharvest head stress and subsequent cold stress at the onset of cold postharvest storage such as delayed sun scald (B & C), which is initiated in the field during periods of intense heat and solar irradiation but does not manifest until fruit are put in cold storage (B - 2 weeks postharvest versus C - 5 months postharvest). Photo credits A) Heidi Hargarten, USDA-ARS TFRL, C&D) Christine McTavish, USDA-ARS TFRL. **ALT text:** Three photographs. A shows several green apples clustered on a leafy tree branch. B and C show a single apple two weeks and five months after harvest, respectively, with a tan/brown discolored patch appearing only in the later image.

The frequency of extreme heat days (> 34 °C) in major U.S. apple-producing regions has doubled over the past four decades, from approximately five to more than 10 days per year ^23^. Since fruit surface temperatures can substantially exceed ambient air temperature ^26,27^, increasing heat exposure elevates the risk of sunburn-related disorders including delayed sunscald in ‘Granny Smith’ (**Figure 1B, C**), which manifests only after cold storage ^26,28–30^. In Washington State, which accounts for roughly half of U.S. apple production, sun and heat overexposure cause average annual losses exceeding 10%, costing growers more than $150 million ^26^. Losses in unprotected orchards in warmer growing regions can be three to four times higher ^26^. Mitigation through netting and evaporative cooling are capital- and labor-intensive ^31,32^, while affected fruit require additional handling ^33^.

Understanding these disorders requires attention to cultivar- and even tissue-specific responses. Studies of postharvest storage, disorder development, and canopy light environment reveal distinct molecular responses among pome fruit cultivars and tissues ^34–38^. For example, sunscald in ‘Granny Smith’ and stain in ‘Fuji’ differ in etiology and molecular underpinning despite both arising from heat and light overexposure and developing symptoms during cold storage ^26,35^. Resolving these responses therefore requires genomic resources that accurately represent the genotype under investigation.

Transcriptomic studies of highly heterozygous, clonally propagated crops like apple have historically relied on a single, often distantly related conspecific reference genome. However, RNA-seq interpretation depends on how well the reference sequence represents the studied genotype ^39,40^. Reads containing non-reference alleles or structural variants may align less efficiently or fail to align, introducing reference bias into differentially expression analyses. In rice, reference choice altered the identity of differentially expressed genes ^40^; in wheat, cultivar introgressions generated spurious expression differences between cultivars ^39^. Genotype-informed human references likewise reduced RNA-seq mapping error two- to three-fold ^41^. In apple, this problem is not hypothetical - candidate qPCR primer-binding sites selected from the ‘Golden Delicious’ (GDDH13) reference genome ^42^ contained problematic polymorphisms in ‘Granny Smith’ and ‘Honeycrisp’, compromising confidence in gene expression measurements ^43^. Substantial divergence between apple haplotypes further limits the ability of collapsed or off-genotype references to represent sequence variation ^44^.

High-quality, genotype-specific reference genomes address these limitations in two complementary ways. First, they support a truer “apples-to-apples” genetic comparison, allowing more reliable identification of biological expression responses and allele-specific variation. In cassava, a haplotype-resolved genome revealed allele-specific expression differences in 20-30% of transcripts per tissue that would otherwise be obscured or biased using a collapsed reference ^45^. In grape, phased assemblies can also improve fine-scale genetic mapping relative to consensus references ^46^. Second, they provide foundations for pangenomes and pantranscriptomes that integrate variation across genotypes, improving mapping accuracy and enabling haplotype-specific transcript quantification ^47^. Together, these resources are increasingly essential for support investigation of cultivar-specific physiology and identification of potential breeding targets and prognostic biomarkers. Expanding genomic resources beyond a single reference cultivar is therefore particularly valuable for crops whose commercial value depends on genetically distinct genotypes.

Here, we present a chromosome-scale, parent-of-origin-resolved genome assembly for ‘Granny Smith’, with assembly BUSCO completeness of 98.6% for both haplotypes and annotation completeness of 98.7% and 98.9% for HapA and HapB, respectively. We then use this resource to ask a broader question: ***how strongly does reference choice influence the biological conclusions drawn from transcriptomic data in a highly heterozygous crop?*** Using RNA-seq data from sun- and shade-exposed ‘Granny Smith’ peel during postharvest cold storage, we compared an earlier-generation non-native reference (GDDH13) ^42^, a modern phased non-native reference (‘WA 38’) ^48^, and the cultivar-matched ‘Granny Smith’ haplotypes presented here. Rather than evaluating mapping performance alone, we followed reference-dependent effects through differential expression, gene-family concordance, and functional enrichment, and compared the resulting biological signals with metabolomic patterns previously measured from the same experimental system. This design allows us to test whether apparently modest differences in reference compatibility translate into meaningful differences in the resolution and interpretation of cultivar-specific physiological responses.

## 2. Results

### 2.1 Deep sequencing reveals a highly heterozygous genome

The three sequencing datasets provided substantial coverage of the ‘Granny Smith’ genome (**Table 1**). PacBio HiFi sequencing generated 73.2 Gb (approximately 113× haploid coverage), while Omni-C sequencing generated 330.6 Gb and Illumina shotgun sequencing generated 107.2 Gb. FastQC profiles were generally consistent with the expected sequence characteristics. Estimated heterozygosity was consistent between the Illumina shotgun and HiFi datasets (1.31– 1.34%), comparable to ‘WA 38’ ^48^ and slightly higher than ‘Honeycrisp’ ^49^. GenomeScope estimated that approximately 69% of the genome is comprised of unique sequences, substantially more than reported for those two cultivars (approximately 52–54%). Full sequencing-quality and genome complexity reports are available through the project GitLab repository (https://gitlab.com/ficklinlab-public/gs-genome-project).

**Table 1:** Sequencing yields from PacBio Long Read (HiFi) and Illumina Short Read (Shotgun and Omni-C). Coverage is estimated based on a haploid genome of 650Mb.

|  | **Long Read** | **Short Read** |  |
|---|---|---|---|
|  | **HiFi** | **Shotgun DNA seq** | **OmniC-Seq** |
| Total read number | 4,692,414 | 711,383,096 | 2,190,184,148 |
| Number of bases (Gb) | 73.2 | 107.2 | 330.6 |
| Sequence length (bp) | 43-47,874 | 151 | 151 |
| Coverage | 113x | 165x | 508x |
| %GC | 38 | 38 | 40 |
| Avg. Heterozygosity | 1.31% | 1.34% | NA |

### 2.2 Parent-of-origin-resolved haplomes are highly contiguous and complete

Three sets of assemblies were generated: contig, scaffold, and final chromosome-level assemblies (assembly statistics provided in **Table S1**). After Omni-C scaffolding, the 17 largest scaffolds in each provisional haplome corresponded to the 17 apple chromosomes. Pedigree-phased SNP markers were used to assign each chromosome to its parental origin, after which ‘Cleopatra’-derived chromosomes were grouped into HapA and ‘Stone Pippin’-derived chromosomes into HapB. The same marker analysis was then used to validate the reorganized haplomes (**Figure 2A**). All chromosomes in HapA and HapB were consistently assigned to a single parent, with no genotype inconsistencies or localized enrichment of missing or incorrectly aligned SNPs. These results support continuous parent-of-origin phase across all 17 chromosome-scale scaffolds of each haplome, in contrast to previously published assemblies that contain intra-chromosomal phase switches (**Figure 2B**).

**Figure 2.**
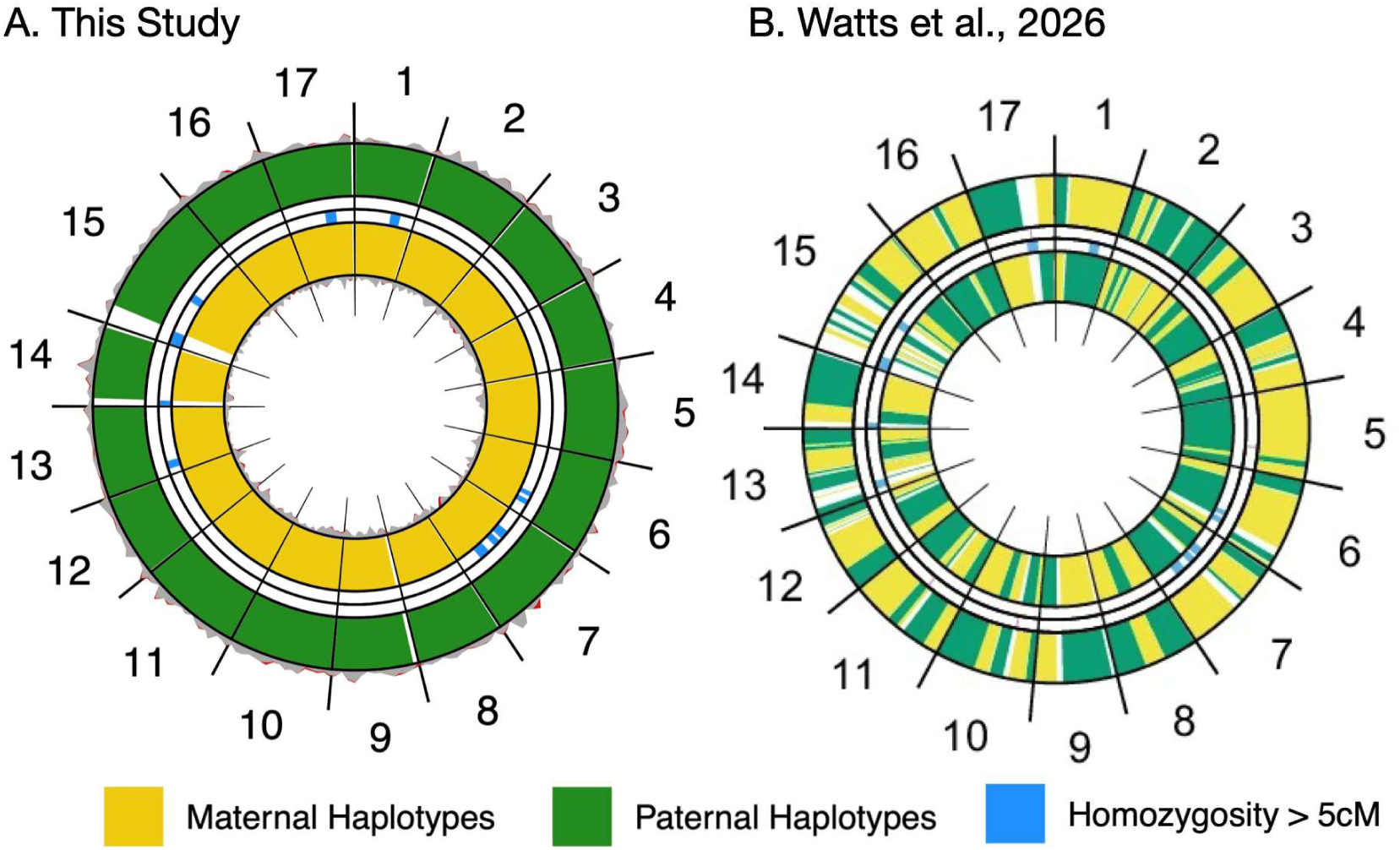
Parental SNP analysis supports chromosome-scale parent-of-origin phasing of the ‘Granny Smith’ genome and reveals contrasting haplotype phasing in the Watts et al. ^61^ **assembly.** Parental haplotype assignments were evaluated for **(A) the ‘Granny Smith’ genome assembled in this study** and **(B) the ‘Granny Smith’ genome reported by Watts et al** ^61^ using reference phased 20K SNP array data obtained from pedigree-based FlexQTL phasing. SNP alleles in HapA (inner wide circle) and HapB (outer wide circle) are assigned as maternal (gold) or paternal (green). Homozygous regions based on SNP array data, particularly at chromosome edges, could not be confidently assigned to either parent and are therefore displayed in white. In the assembly generated in this study (A), all chromosomes within each haplome are consistently assigned to a single parent, with HapA maternal and HapB paternal, supporting chromosome-scale parent-of-origin phasing. In contrast, the Watts et al. ^61^ assembly (B) shows frequent switches between maternal and paternal assignments within chromosomes in both haplomes, revealing multiple intra-chromosomal phase switches. Homozygous regions >5 cM are displayed in blue in the inner thin circle. Genotype inconsistencies between each assembly and the reference SNP array data would be displayed in red in the outer thin circle. Grey or red peaks in the innermost and outermost circles indicate the proportion of SNPs that are not assigned to a single region or are assigned to the incorrect chromosome, respectively. ALT TEXT: Side-by-side circular plots of the 17 apple chromosomes comparing parental haplotype assignments in two ‘Granny Smith’ genome assemblies. In panel A, representing this study, the outer haplotype ring is consistently green (paternal) and the inner haplotype ring is consistently gold (maternal) across nearly all chromosomes, with small white regions at some chromosome edges and scattered blue marks indicating homozygosity greater than 5 cM. In panel B, representing the Watts et al. (2026) assembly, both the outer and inner haplotype rings contain numerous alternating green and gold segments within chromosomes, indicating frequent switches between paternal and maternal haplotype assignments.

The final HapA assembly comprised 17 chromosome-scale scaffolds and eight unplaced scaffolds, totaling 651.0 Mb, whereas HapB comprised 17 chromosome-scale scaffolds and six unplaced scaffolds, totaling 648.0 Mb (**Table 2**). Scaffold N50 values were 36.5 and 37.7 Mb, respectively, and both assemblies contained 98.6% complete BUSCOs. LTR Assembly Index (LAI) scores of 24.80 for HapA and 23.45 for HapB placed both assemblies in the gold-quality category for repetitive and intergenic regions ^50^.

**Table 2.** Summary statistics of the final ‘Granny Smith’ Hap A and B assembly and annotation. Includes a selection of previously published *Malus domestica* genomes for comparison. Some genome statistics were re-run for more accurate comparison (see Zhang et al ^48^).

|  | **Granny Smith** |  | **Granny Smith** |  | **WA 38** |  | **Honeycrisp** |  | **Golden Delicious T2T** |  | **Fuji** | **GDDH13** |
|---|---|---|---|---|---|---|---|---|---|---|---|---|
|  | **HapA** | **HapB** | **Hap1** | **Hap2** | **HapA** | **HapB** | **HapA** | **HapB** | **Hap1** | **Hap2** |  |  |
| n Scaffolds | 17 (+8) | 17 (+6) | 17 | 17 | 17 | 17 | 473 | 215 | 17 | 17 | 1,358 | 1,081 |
| Haploid Genome size (Mb) | 651 | 648 | 653 | 645 | 645 | 651 | 674 | 660 | 651 | 645 | 736.9 | 709.6 |
| N50 (Mb) | 36.5 | 37.7 | 37.7 | 36.5 | 36.1 | 37.2 | 31.6 | 32.8 | 37.7 | 36.3 | 36.8 | 5.5 |
| L50 | 8 | 8 | 8 | 8 | 8 | 8 | 8 | 8 | 8 | 8 | 9 | NA |
| n Protein Coding Genes | 54,474 | 58,139 | 54,320 | 53,383 | 53,028 | 54,235 | 47,563 | 48,655 | 44,375 | 44,304 | 49,972 | 45,116 |
| Complete BUSCO (%) Assembly | 98.6 | 98.6 | 98.3 | 98.0 | 98.7 | 98.7 | 98.6 | 98.7 | 98.7 | 98.6 | 98.8 | 98 |
| Complete BUSCO (%) Annotation | 98.7 | 98.9 | 95.6 | 95.3 | 98.5 | 98.4 | 96.8 | 97.4 | 96.4 | 97.0 | 97.2 | 96.1 |
| n Orthogroups (26Gv2.0) | 10,464 | 10,443 | 10,325 | 10,271 | 10,494 | 10,511 | 10,350 | 10,366 | 9,696 | 9,703 | 10,243 | 10,117 |
| Reference | This study |  | Watts et al., 2026 |  | Zhang et al., 2024 |  | Khan et al. 2022 |  | Su et al., 2024 |  | Li et al. 2024 | Daccord et al. 2017 |

Among published apple genomes, ‘Granny Smith’ most closely matched the chromosome-scale contiguity of ‘WA 38’ ^48^ and ‘Golden Delicious’ T2T ^51^, with all 17 chromosomes represented and few unplaced scaffolds (**Table 2**). The ‘Granny Smith’ haplomes were less fragmented than several other cultivar assemblies, including ‘Honeycrisp’, while retaining similarly high assembly BUSCO completeness (slightly higher than the ‘Granny Smith’ assemblies released by Watts et al ^61^). These comparisons establish ‘Granny Smith’ as a highly contiguous, parentally resolved reference genome.

### 2.3 Repeat annotation reveals broadly conserved haplome landscapes

Telomeric repeats were detected at one or both ends of all 17 chromosome-scale scaffolds in each haplome, including both ends of 11 chromosomes (**Table S2**). The predominant motif, AAACCCT and its reverse complement AGGGTTT, was identical to that reported for ‘WA 38’ ^48^ and consistent with the canonical flowering-plant telomeric repeat ^52^.

RepeatMasker identified repeats across 59.65% of HapA and 59.07% of HapB, with LTR retrotransposons comprising approximately 40% of each assembly (**Table S3**). Repeat composition was highly similar between haplomes, although RTE-family LINEs were more abundant in HapA (**Table 3**). The two haplomes also showed similar repeat-divergence profiles, characterized by a concentration of low-divergence LTR elements and a second increase near 9% divergence, consistent with recent and older periods of LTR accumulation (**Figure S1**).

**Table 3.** Repeat coverage of ‘Granny Smith’ Hap A and Hap B. , as well as ‘WA 38’ and ‘Honeycrisp’ haplomes for comparison. RTE-family LINE transposon information is not available in **‘**WA 38**’** and **‘**Honeycrisp**’.**

| **Class** |  | **'Granny Smith' (%)** |  | **'WA 38' (%)** |  | **'Honeycrisp' (%)** |  |
|---|---|---|---|---|---|---|---|
|  |  | **HapA** | **HapB** | **HapA** | **HapB** | **HapA** | **HapB** |
| **LINE** | L1 | 0.36 | 0.54 | 0.00 | 0.00 | 0.00 | 0.00 |
|  | RTE | 3.56 | 0.01 | N/A | N/A | N/A | N/A |
| **LTR** | Copia | 6.66 | 7.29 | 9.37 | 10.22 | 9.73 | 9.60 |
|  | Gypsy | 12.57 | 12.02 | 17.19 | 18.32 | 20.29 | 17.80 |
|  | unknown | 20.14 | 22.32 | 16.37 | 14.52 | 14.89 | 16.86 |
|  | **Total** | **39.37** | **41.63** | **42.93** | **43.06** | **44.91** | **44.26** |
| **TIR** | CACTA | 1.49 | 1.50 | 1.94 | 2.15 | 2.21 | 1.95 |
|  | Mutator | 2.52 | 2.69 | 3.96 | 4.18 | 4.16 | 4.25 |
|  | PIF_Harbinger | 3.55 | 3.35 | 2.40 | 2.52 | 2.43 | 2.60 |
|  | Tc1_Mariner | 0.09 | 0.07 | 0.16 | 0.24 | 0.15 | 0.27 |
|  | hAT | 2.13 | 2.19 | 2.15 | 2.37 | 2.30 | 2.31 |
|  | polinton | 0.00 | 0.00 | 0.00 | 0.00 | 0.00 | 0.01 |
|  | **Total** | **9.78** | **9.80** | **10.61** | **11.46** | **11.25** | **11.39** |
| **nonLTR** | pararetrovirus | 0.24 | 0.17 | 0.00 | 0.00 | 0.00 | 0.00 |
|  | LINE_element | 0.00 | 0.00 | 0.14 | 0.16 | 0.18 | 0.17 |
|  | unknown | 0.00 | 0.00 | 0.11 | 0.09 | 0.09 | 0.18 |
|  | **Total** | **0.24** | **0.17** | **0.25** | **0.25** | **0.27** | **0.35** |
| **nonTIR** | helitron | 1.52 | 2.10 | 3.41 | 2.20 | 2.95 | 3.18 |
| **Other repeat Region** |  | 1.46 | 0.35 | 1.52 | 1.74 | 2.91 | 2.78 |
| RepeatMasker |  | 3.36 | 4.47 | 3.98 | 3.99 | NA | NA |
| **Total** |  | **59.65** | **59.07** | **62.71** | **62.71** | **62.43** | **61.97** |
| Reference |  | This Manuscript |  | Zhang et al., 2024 |  | Khan et al., 2022 |  |

Total repeat coverage was slightly lower than in ‘WA 38’ ^48^ and ‘Honeycrisp’ ^49^, principally reflecting lower Copia, Gypsy, and TIR content, whereas unknown LTRs represented a larger fraction of the ‘Granny Smith’ assemblies (**Table 3**). Despite these cultivar-level differences, the close agreement between HapA and HapB indicates broadly conserved repeat landscapes across the two parental haplotypes.

### 2.4 Gene-space and functional annotations are highly complete

The final annotations contained 54,474 protein-coding genes in HapA and 58,139 in HapB and retained 98.7% and 98.9% complete BUSCOs, respectively (**Table 2**). Although HapB contained 3,665 more genes, the two annotations had similar transcript and coding-sequence characteristics (**Table S4**). Statistics for intermediate annotations are provided in **Table S4**.

The final ‘Granny Smith’ annotations contained more protein-coding genes than the other apple genomes in **Table 2**, with HapB having the largest gene set and HapA slightly exceeding both ‘WA 38’ haplomes. Despite these higher gene counts, annotation BUSCO completeness was also highest for the two ‘Granny Smith’ haplomes and was broadly comparable to ‘WA 38’. Reanalysis of the published Watts et al ^61^ ‘Granny Smith’ protein sets using the same eudicots_odb10 lineage recovered 95.6% and 95.3% complete BUSCOs for haplotypes 1 and 2, respectively, compared with 98.7% and 98.9% for HapA and HapB in this study. Together, these results indicate that the ‘Granny Smith’ annotations presented here provide highly complete representations of gene space.

At least one InterProScan or Enzyme Commission annotation was assigned to 92.1% of HapA genes and 94.0% of HapB genes, whereas 48.7% and 46.1% having high confidence hits in the Uniprot/Swiss-prot database^53^ (**File S1**). HapA had slightly higher proportions of genes with InterPro and Gene Ontology assignments, whereas Enzyme Commission annotation counts were nearly identical between haplomes. Overall functional-annotation coverage was comparable to, and slightly higher than, that reported for ‘WA 38’ ^48^, providing functional information for most genes in both ‘Granny Smith’ haplomes.

### 2.5 Gene-family content and chromosome organization are broadly conserved

#### 2.5.1 Most gene families are shared between haplomes

Protein-coding genes from the two ‘Granny Smith’ haplomes generated in this study were classified into PlantTribes2 26Gv2.0 orthogroups and compared with published *Malus* and *Pyrus* annotations, including the ‘Granny Smith’ haplotypes reported by Watts et al ^61^. Our HapA and HapB annotations, designated Md.GS.A and Md.GS.B, represented 10,464 and 10,443 orthogroups, respectively (**Table 2**), with 10,263 shared between them. In this pairwise comparison, 201 orthogroups were detected only in HapA and 180 only in HapB. Most were also conserved across other apple genomes: 8,275 orthogroups were present in all included *M. domestica* annotations (**Figure 3B**), and 7,790 remained shared when wild apple and *P. communis* ‘d’Anjou’ outgroups were included (**Figure S2**). Notably, the 312 orthogroups absent from both ‘Golden Delicious’ T2T haplomes but present in every other *M. domestica* annotation formed the second-largest intersection in the UpSet analysis, suggesting that this pattern may reflect systematic annotation differences rather than true gene loss.

**Figure 3.**
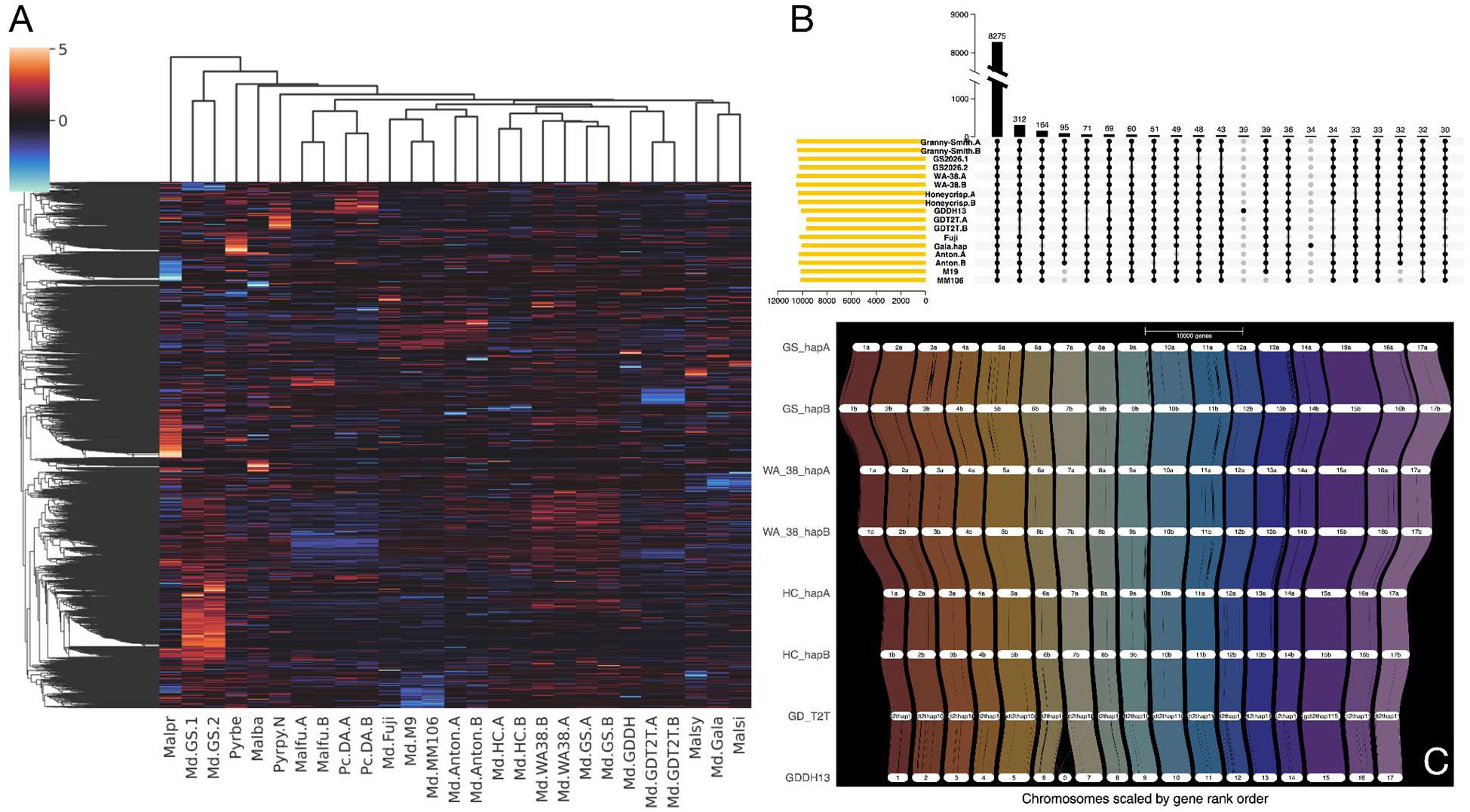
Comparative gene-family content and chromosome-scale synteny across apple and related Maleae genomes. **(A)** Hierarchical clustering of standardized gene counts for core orthogroups (CROGs) across the included *Malus* and *Pyrus* annotations. Columns represent genome or haplome annotations and rows represent CROGs; warm and cool colors indicate gene counts above and below the cross-annotation mean, respectively. **(B)** UpSet plot showing orthogroup intersections among the included *M. domestica* annotations. Horizontal bars indicate the total number of orthogroups in each annotation, vertical bars indicate intersection sizes, and connected black dots identify the annotations comprising each intersection. The largest intersections are shown. **(C)** GENESPACE comparison of chromosome-scale synteny among ‘Granny Smith’ (GS), ‘WA 38’, ‘Honeycrisp’ (HC), ‘Golden Delicious’ T2T (GD_T2T), and GDDH13. Chromosomes are scaled by gene rank order, and colored ribbons connect syntenic regions between genomes. The predominance of vertical ribbons demonstrates extensive one-to-one chromosome correspondence, whereas localized crossings indicate reverse-oriented or rearranged regions. In panel A, Md denotes *M. domestica*; Malba, *M. baccata*; Malfu, *M. fusca*; Malpr, *M. prunifolia*; Malsi, *M. sieversii*; Malsy, *M. sylvestris*; Pyrbe, *Pyrus betulifolia*; Pyrpy.N, *P. pyrifolia* ‘Nijisseiki’; and P.c.DA, *P. communis* ‘d’Anjou’. Md.GS.A and Md.GS.B represent the ‘Granny Smith’ HapA and HapB annotations generated in this study, whereas Md.GS.1 and Md.GS.2 represent the ‘Granny Smith’ haplotype 1 and haplotype 2 annotations published by Watts et al ^61^. **ALT TEXT:** Three panels. A is a clustered heatmap showing gene-count variation across Malus and Pyrus genomes, with related species grouping together in the dendrogram. B is an UpSet plot showing that most orthogroups are shared across all seventeen apple annotations, with only small subsets unique to individual genomes. C shows chromosome-scale synteny ribbons linking seventeen chromosomes across five apple genomes, mostly straight and vertical with a few crossing regions.

To examine variation in annotated gene counts within conserved families, we compared 9,656 CROGs across 27 Malus and Pyrus annotations (**Figure 3A**). Md.GS.A and Md.GS.B clustered together within a broader group containing ‘WA 38’ and ‘Honeycrisp’. The ‘Golden Delicious’ T2T haplomes formed a pair that grouped with GDDH13. In contrast, the two ‘Granny Smith’ haplomes from Watts et al, Md.GS.1 and Md.GS.2, formed a distinct pair separated from the principal cluster containing most other *Malus* annotations, including our ‘Granny Smith’ genome. Both Watts et al ^61^ annotations displayed conspicuous regions of elevated relative gene counts and broader z-score distributions (**Figure S3**): their interquartile ranges were -0.24 to 1.30 and -0.26 to 1.28, compared with -0.25 to 0.68 for Md.GS.A and -0.28 to 0.59 for Md.GS.B. Of the full CROG set, 9,603 and 9,600 were represented in Md.GS.A and Md.GS.B, compared with 9,489 and 9,466 in Md.GS.1 and Md.GS.2, respectively. Furthermore, the previously observed “publication bias” remained evident, with annotations produced by the same study or similar workflows tending to cluster together ^48^.

Despite their overall similarity, Md.GS.A and Md.GS.B differed substantially in a small subset of CROGs. Counts were identical in 6,205 CROGs and differed by no more than two genes in 9,191 of 9,656 CROGs (95.2%). Four CROGs contained more than 50 additional genes in HapB and five contained more than 50 additional genes in HapA (**Table 4**). The largest difference occurred in the RING/U-box family OG2281, with 283 genes in HapA and 2,954 in HapB. Several HapB-enriched outliers also differed substantially in predicted protein length: mean proteins for OG2281, OG6885, and OG5581 were considerably shorter in HapB than in HapA. These results identify specific gene families that warrant further investigation to distinguish genuine haplotype-specific expansions from differences introduced during gene-model prediction.

**Table 4.** CROG outlier summary.

| OG | Other annotations: median (range) | HapA genes | HapB genes | Mean protein length, HapA (aa) | Mean protein length, HapB (aa) | Interpretation |
|---|---|---|---|---|---|---|
| OG2281 | 20 (2–1,006) | 283 | 2,954 | 983 | 361 | Highly variable; HapB is an extreme outlier |
| OG6885 | 3 (2–40) | 3 | 83 | 385 | 125 | HapA is typical; HapB exceeds all others |
| OG5581 | 3 (2–7) | 3 | 69 | 342 | 162 | HapA is typical; HapB is an extreme outlier |
| OG502 | 48 (3–136) | 99 | 150 | 258 | 279 | Broad variation; both are near the upper range |
| OG26 | 70 (2–483) | 476 | 224 | 335 | 309 | Broad variation; HapA is near the maximum |
| OG64 | 45 (11–106) | 229 | 61 | 439 | 489 | HapB is typical; HapA exceeds all others |
| OG2368 | 5 (0–99) | 83 | 9 | 241 | 199 | Variable; both remain within the observed range |
| OG423 | 26 (1–89) | 111 | 43 | 195 | 183 | HapA is moderately above the other annotations |
| OG6 | 310 (149–407) | 413 | 361 | 718 | 778 | Generally high-copy family; both are within or near the range |

#### 2.5.2 Chromosome organization is strongly conserved across Maleae

The two ‘Granny Smith’ haplomes shared 37,799 syntenic gene anchors across 31 regions, all connecting corresponding chromosomes (**Figure 3C**). Most regions were collinear, with seven localized reverse-oriented regions; the largest spanned approximately 6.6 Mb on chromosome 3. Comparisons with other domesticated apple genomes likewise showed one-to-one chromosome correspondence.

Chromosome-scale synteny was similarly conserved across representative wild *Malus* species and *Pyrus communis* ‘d’Anjou’ (**Figure S4**). Nearly all anchors connected corresponding chromosomes, including 25,851 of 25,974 anchors between ‘Granny Smith’ HapA and ‘d’Anjou’. These results indicate that the 17-chromosome organization and overall gene order are strongly conserved across the examined Maleae genomes, with structural differences concentrated in localized regions.

### 2.6 The ‘Granny Smith’ organellar genome assembly and annotation

The ‘Granny Smith’ chloroplast genome was 160,069 bp and displayed the typical large single-copy, small single-copy, and paired inverted-repeat organization, with genes supporting photosynthesis, transcription, and translation (**Figure S5**). The 232,437-bp mitochondrial assembly contained genes encoding respiratory-chain complexes, ATP synthase, ribosomal proteins, rRNAs, and tRNAs (**Figure S6**). Compared with the gene-dense chloroplast genome, mitochondrial genes were more sparsely distributed across the assembly, with substantial intervening noncoding regions.

### 2.7 Comparative transcriptomics reveals reference-dependent biological inference

#### 2.7.1 RNA-seq mapping rates are modestly improved with cultivar-matched references

RNA-seq data from sun- and shade-exposed ‘Granny Smith’ peel during postharvest cold storage were mapped to four different assemblies, 1) GDDH13 (earlier generation, non-native cultivar), 2) ‘WA 38’-HapA (new generation, non-native cultivar), 3) ‘Granny Smith’-HapA (GS-HapA; new generation, native cultivar), and 4) GS-HapB (new generation, native cultivar). Overall mapping rates were high, with a small number of samples showing consistently low rates across all four experiments. Median mapping rates are reported here; full details are provided in **File S2, Table S5**. When mapping ‘Granny Smith’ RNA-Seq reads to the matching cultivar haplomes, median uniquely mapped read rates were approximately one percentage point higher compared to mapping to either off-cultivar genome (GDDH13=93.2%; ‘WA 38’=93.2%; GS-HapA=94.3%; GS-HapB=94.1%). Similarly, the number of unmapped reads was higher when mapping to the off-cultivar genome (GDDH13=1.9%, ‘WA 38’-HapA=2.7%, GS-HapA=1.1%, GS-HapB=1.1%). Thus, cultivar matching increased unique mapping by only ∼1% while reducing unmapped reads. We next asked whether these comparatively modest differences in alignment performance propagated into larger differences in downstream biological inference.

#### 2.7.2 Differentially expressed genes show broadly conserved expression trajectories across references

We focused on differential expression at key time points based on McTavish et al ^35^ findings regarding postharvest metabolomic onset of delayed sun scald; 1) the difference between sun and shade exposed tissue at harvest (SunVsShd_T0), 2) the effect of time in storage (T0 to 2-weeks cold storage) on each side of the fruit (Sun_2wkVsT0 and Shd_2wkVsT0), and 3) the interaction effect of sun/shade exposure and time in storage at 2-weeks postharvest (SunVsShd_2wk) (**Table 5**).

**Table 5.**
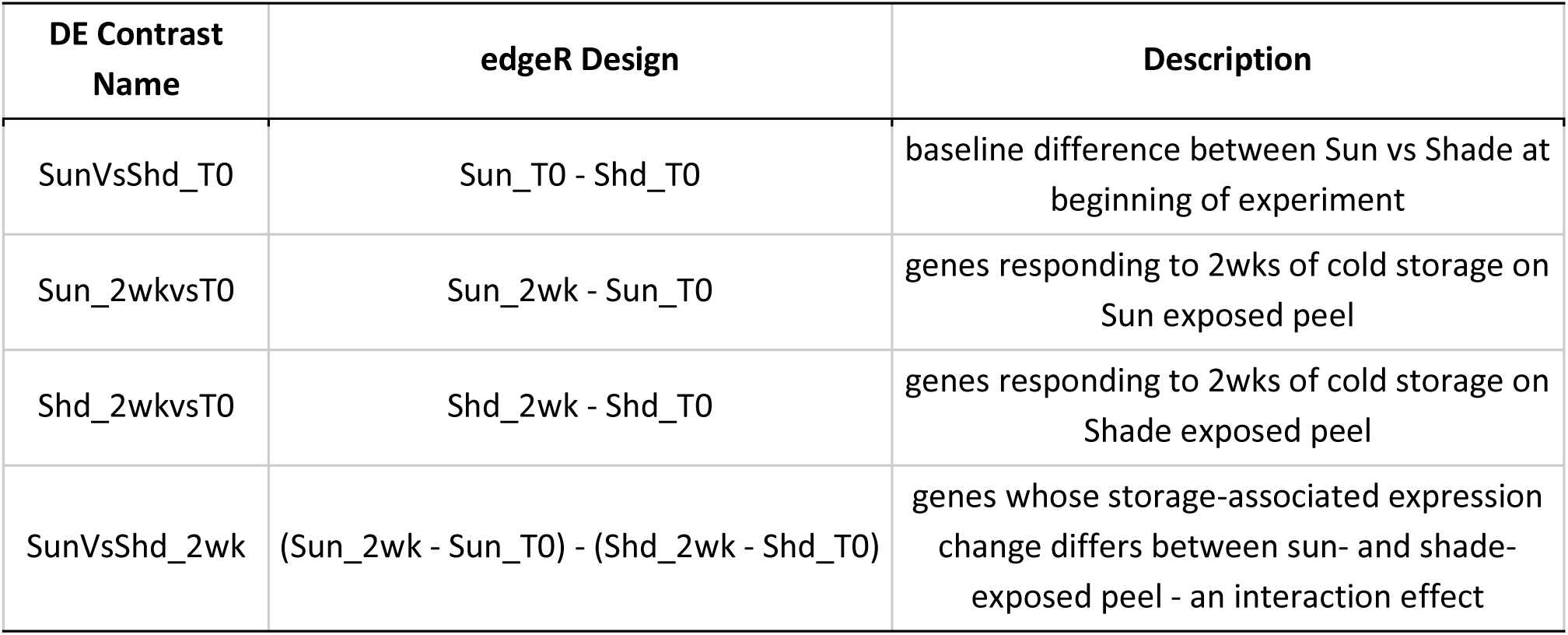
Differential Expression (DE) Contrasts selected for mapping experiment case study. DE Contrast Name is used throughout the manuscript to reference the DE design being referenced. edgeR Design is the DE design to make the desired DE contrasts. Description describes what each DE contrast represents.

| DE Contrast Name | edgeR Design | Description |
|---|---|---|
| SunVsShd_T0 | Sun_T0 - Shd_T0 | baseline difference between Sun vs Shade at beginning of experiment |
| Sun_2wkvsT0 | Sun_2wk - Sun_T0 | genes responding to 2wks of cold storage on Sun exposed peel |
| Shd_2wkvsT0 | Shd_2wk - Shd_T0 | genes responding to 2wks of cold storage on Shade exposed peel |
| SunVsShd_2wk | (Sun_2wk - Sun_T0) - (Shd_2wk - Shd_T0) | genes whose storage-associated expression change differs between sun- and shade-exposed peel - an interaction effect |

Clusters of DEGs with similar expression patterns were synchronized across mapping experiments to facilitate direct comparison (**Tables S6-S7, File S3**). Overall, expression profiles of DEGs had similar cluster profiles across all genomes throughout the experiment (**Figure 4**). The DE experiment SunVsShd_2wk was the only contrast that had different numbers of clusters identified across selected genomes. In this DE contrast, GDDH13 DE down regulated genes were assigned to 4 different clusters while the other three genomes had 2 clusters, and GS-HapA DE up regulated genes were assigned to 2 different clusters while the other three genomes had 3 clusters. Correlation between synchronized cluster centroid expression profiles (**Table S6**) were highest between GS-HapA and GS-HapB (mean r = 0.980), followed closely by GS-HapA and ‘WA 38’ (mean r = 0.978), with GS-HapA vs GDDH13 having the lowest correlation (mean r = 0.944). Thus, broad expression trajectories were highly conserved across references despite differences in individual DEG membership and cluster structure.

**Figure 4.**
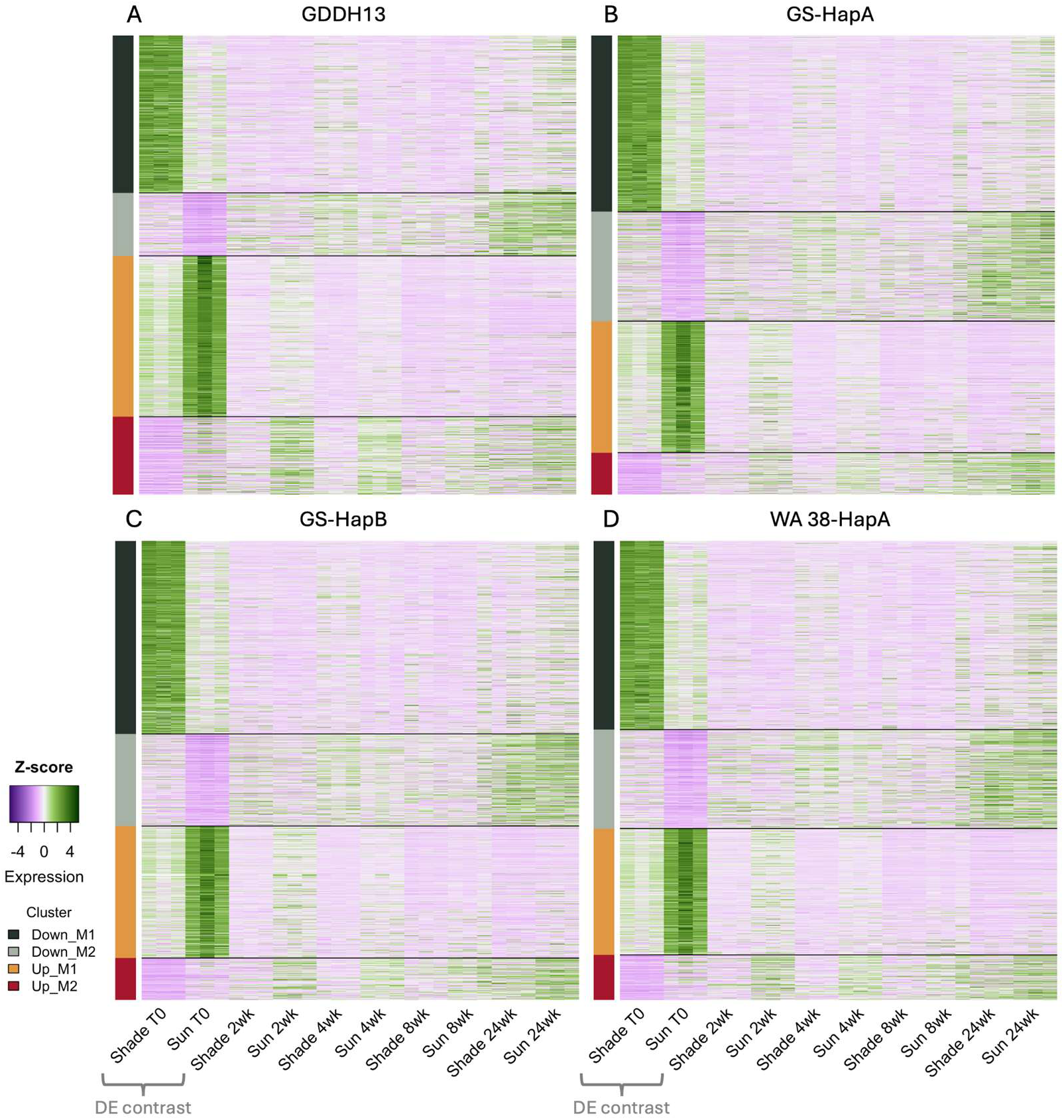
Expression trajectories of genes differentially expressed between sun- and shade-exposed peel at harvest across four mapping experiments. DE contrast is between the sun exposed side of ‘Granny Smith’ apple peel versus the shade protected side - Up and Down regulation designations are with reference to the sun side, ie up/down regulated on the sun side versus the shade side. Synchronized clusters of genes with similar expression patterns highlight differences in cluster membership and behavior throughout the postharvest storage period (at harvest through 24 weeks cold storage) both within and across genomes. Cluster size and membership are detailed in TableS7 and FileS3. ALT TEXT: Four heatmaps, one per genome assembly, showing gene expression across sun and shade samples at five postharvest storage timepoints. In all four, expression differences between sun and shade are strongest at time zero and diminish over storage, with a consistent pattern across the four gene clusters.

#### 2.7.3 Functional Enrichment reveals core functions and notable disparity among mapping experiments

In contrast to the broadly conserved expression trajectories, functional enrichment profiles diverged substantially among references. All four mapping experiments recovered a core set of shared biological responses, but most enriched terms were not shared across all references (**Figure 5**; **Files S4-S5**). At harvest, chloroplast thylakoid membrane GO term (GO: 0009535) was enriched among sun-upregulated genes in all genomes (SunVsShd_T0 - **Files S4.1, S4.5, S4.9, S4.13**; GS-HapA: 17 genes, FDR = 5.4×10⁻¹¹; GDDH13: 5 genes, FDR = 9.1×10⁻⁵; WA 38: 4 genes, FDR = 1.3×10⁻⁴), with the largest number of contributing genes in GS-HapA, consistent with more genes in total being annotated with this GO term in the GS-HapA genome compared to the GDDH13 and WA 38 genomes (160 vs. 14 and 26, respectively). This harvest-time difference is then succeeded by a much larger, general decline during storage: photosystem/chloroplast terms were strongly enriched among storage-down regulated genes on both sun- and shade-exposed sides (Sun_2wkVsT0 & Shd_2wkVsT0 - **Files S4.2, S4.3, S4.6, S4.7, S4.10, S4.11, S4.14, S4.15**; FDR <10⁻⁸ in multiple contrasts), and by two weeks of postharvest cold storage this photosynthetic signal was no longer significantly enriched in the SunVsShd_2wk interaction contrast (**Files S4.4, S4.8, S4.12, S4.16**).

**Figure 5.**
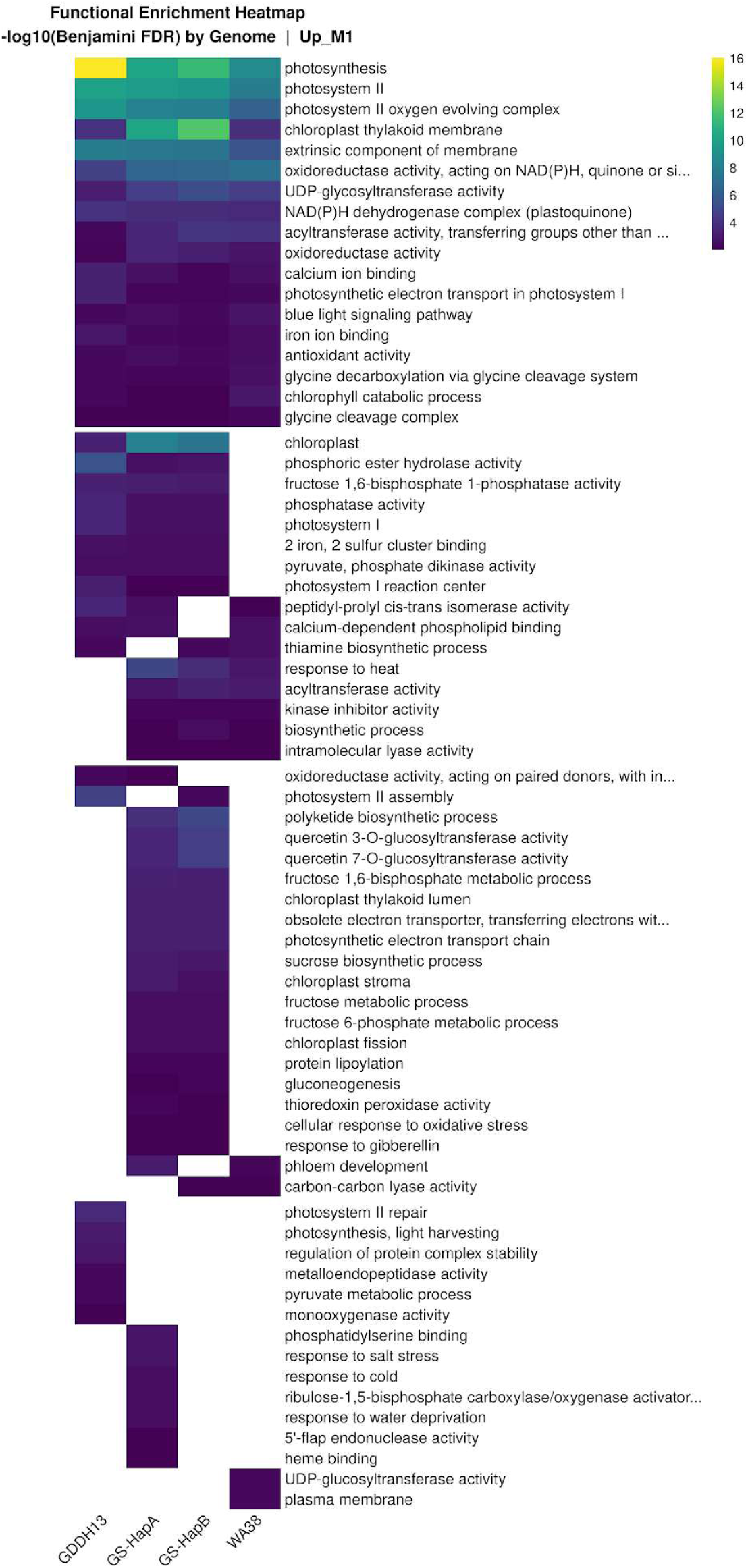
At harvest Functional Enrichment (FE) for up-regulated genes on the sun exposed peel in expression cluster 1 across four mapping experiments. A core group of shared enriched functions is apparent and terms for photosystem, chlorophyll, and light response associated processes show clear cross genome functional corroboration. However, the bulk of enriched terms are not shared across all genomes. Plot cells are scaled by -log10 Benjamini corrected false discovery rate (FDR) within a genome. Enriched functions listed are descriptive GO terms. Full FE results can be found in FileS4 and comparative FE heatmaps for the additional FE contrasts of interest in FileS5. ALT TEXT: Heatmap of gene ontology enrichment significance across four genome assemblies. A shared core of photosynthesis and chlorophyll related terms near the top shows strong enrichment in all four genomes, while the majority of listed terms below are enriched in only one or a few genomes, with several cells blank.

The size of this core group of functions was typically dwarfed by enriched terms that were unique to GS-HapA and/or GS-HapB, with the magnitude of the difference varying by DE contrast and expression clusters (**Files S4-S5**). Notably, GDDH13 had the fewest enriched terms across all comparisons (**File S4**), and its clusters often differ functionally from those identified using ‘Granny Smith’ and ‘WA 38’ (ex. **Files S5.6 & S5.10**). When testing the interaction effect between the sun side and shade side after 2-weeks of cold storage, the DE expression clusters and their functions did not appear to corroborate each other across the three genomes (**Files S5.13-S5.18**).

Genome-dependent differences included functions associated with fruit response to high light and heat stress and postharvest cold storage. For example, at harvest, oxidative-stress enrichment (glutathione/thioredoxin peroxidase, response to oxidative stress) was detected only in GS-HapA (**Files S4.1, S4.5, S4.9, S4.13**). Flavonoid biosynthesis (affected by UV light exposure ^54^ and influences fruit color/flavor ^55^) was enriched among storage-upregulated genes in sun-exposed ‘WA 38’ (Sun_2wkVsT0; 3/15 genes, FDR = 6.0×10⁻³, **File S4.14**) and shade-exposed GS-HapA (Shd_2wkVsT0; 3/6 genes, FDR = 7.8×10⁻³, **FileS 4.6**), but not in either sun-vs-shade contrast (SunVsShd_T0, **File S4.5 & File S4.13**; SunVsShd_2wk, **File S4.8 & File S4.16**). Sterol biosynthesis (associated with postharvest chilling disorders ^56^) was enriched among sun-downregulated genes at harvest in GS and ‘WA 38’ (**File S4.5, File S4.9, & File S4.13)**; after storage, enrichment was restricted to GS and associated with sun-upregulated genes (**File S4.8 & File S4.12**). GDDH13 showed no sterol-pathway enrichment (**File S4.1-File S4.4**). Isoprenoid/diterpenoid biosynthesis (associated with postharvest chilling disorders ^57,58^) was enriched among sun-downregulated genes at harvest in GDDH13 and both GS haplomes (**Files S4.1, S4.5, S4.9, S4.13**). However, the same term was enriched among sun-upregulated in all genomes in both Sun_2wkVsT0 (**Files S4.2, S4.6, S4.10, S4.14**) and SunVsShd_2wk (**Files S4.4, S4.8, S4.12, S4.16)**. Finally, lipid transfer activity (membrane integrity under irradiation and oxidative stress ^26^) was enriched among storage-upregulated genes in sun-exposed peel in GDDH13 and the GS haplomes (Sun_2wkVsT0; **Files S4.2, S4.6, S4.10, S4.14),** whereas cutin biosynthesis (associated with cell protection against water loss ^59^) was enriched only in the GS haplomes in the 2-week postharvest interaction effect between sun and shade exposed peel. Several of these reference-dependent enrichments were also consistent with metabolomic patterns previously measured in the same experimental system ^35^, particularly those associated with oxidative stress, sterol metabolism, and cuticle biology.

Enrichment profiles also differed between GS-HapA and GS-HapB. Concordance for GO terms enriched in the same DE direction ranged from Jaccard indices of 0.37 to 0.73, with the highest agreement at harvest (**Table S8**). In Sun_2wkVsT0, one of the best-powered contrasts, concordance was lower for upregulated than downregulated terms (0.37 versus 0.61), suggesting that enrichment differences were not solely attributable to data availability. In SunVsShd_2wk, GS-HapB had an additional upregulated expression cluster (Up_M3) and more haplotype-exclusive upregulated GO terms than GS-HapA (17 versus 4; **Tables S6 and S8**).

Haplotype-specific enrichment also differed in the biological processes recovered. GS-HapA-specific functions included secondary metabolism, hormone responses, and detailed chloroplast processes. In Shd_2wkVsT0, GS-HapA uniquely recovered response to ethylene and auxin, jasmonic acid biosynthesis, flavonoid biosynthesis, and chalcone isomerase activity. In Sun_2wkVsT0, chloroplast organization, chloroplast nucleoid, and photosystem II stabilization were enriched only in GS-HapA, providing additional functional detail beyond the broad chloroplast response detected across references (**Files S4–S5**). By contrast, GS-HapB-specific enrichment during storage included fatty-acid and triglyceride biosynthesis, acetyl-CoA biosynthesis, and diacylglycerol O-acyltransferase activity. In the SunVsShd_2wk interaction, GS-HapB additionally recovered functions associated with abscisic acid, jasmonic acid, and salicylic acid metabolism and methyl-ester hydrolysis.

Several functions recovered using GS-HapB were also detected with one or both non-native references but not GS-HapA. These included fatty-acid biosynthesis during sun-side storage, fatty-acid β-oxidation during shade-side storage, and photosystem II assembly and β-amylase activity at harvest. Lipid oxidation and thiamine biosynthesis were enriched at harvest in GS-HapB, GDDH13, and ‘WA 38’ but not GS-HapA, whereas xyloglucan metabolism and xyloglucosyl transferase activity during shade-side storage were recovered using GDDH13 and ‘WA 38’ with responses consistent in direction with GS-HapB. Thus, neither GS haplome alone captured every enriched response detected across references.

Together, these results show that reference choice affects the functional profiles inferred from differential expression, with particularly strong differences involving GDDH13 and the subtle SunVsShd_2wk interaction. We therefore compared differentially expressed orthogroups across references to quantify gene-family-level agreement and identify potential sources of reference-dependent disagreement.

##### 2.7.3.1 Gene Family Functional Profiles - Do the genomes agree?

First, we assessed agreement among GDDH13, GS, and ‘WA 38’ in the orthogroup (OG) profiles of DEGs across experiments of interest. Concordance was genome dependent: GS and ‘WA 38’, the two modern, phased genomes, agreed more closely with each other than either did with GDDH13. We evaluated concordance using binary Jaccard test, which measures overlap in differentially expressed OGs, and weighted Tanimoto test, which also accounts for response direction and magnitude. In SunVsShd_T0, GS-HapA/HapB and ‘WA 38’ show the highest cross genome concordance on both metrics (Jaccard = 0.819/0.817, Tanimoto = 0.710/0.681), while comparisons involving GDDH13 yielded consistently lower scores, a pattern that holds up across all contrasts (**Table 6**). Reference dependence was greatest for the SunVsShd_2wk interaction. Jaccard similarity between GS and ‘WA 38’ remained 0.74-0.76, whereas comparisons involving GDDH13 fell to approximately 0.19-0.20; only 78-79 DE orthogroups were shared in comparisons involving GDDH13, compared with >330 among the modern references. Despite high overall concordance, GS-HapA/HapB and ‘WA 38’ consistently has the largest Jaccard– Tanimoto gap (ex. SunVsShd_T0 = 0.109/0.136), indicating stronger agreement in OG membership than in logFC direction and magnitude. Interestingly, GS-HapA and GS-HapB had a higher gap between similarity membership and magnitude/direction than comparisons with GDDH13. These gaps seem to be driven by a small number of orthogroups. For example, between GS-HapA and ‘WA 38’ at harvest, only 7 of 886 shared orthogroups show opposite-sign representative logFC values between the two genomes (**Table S9**). The most extreme case in this example is OG 3, whose representative paralog had a logFC of -10.0 in GS-HapA and +1.15 in ‘WA 38’. Despite differences in detection and effect magnitude, direction was highly concordant among shared DE orthogroups, with 94–99% showing the same sign across pairwise comparisons. Given the generally high orthogroup-level similarity between GS-HapA and GS-HapB (Jaccard = 0.873), we present gene family level analyses below with GS-HapA only.

**Table 6.** Similarity Scores across six pairwise genome comparisons for the four DE contrasts of interest. Genome A/B: The two genomes in the pairwise comparison; Jaccard Score: Binary detection agreement — of orthogroups significant in either genome, the fraction of OGs significant in both; Tanimoto Score: Magnitude- and sign-aware similarity on the representative-logFC vectors - penalizes both absence and sign/magnitude disagreement; OGs Shared: n orthogroups where both genomes independently called at least one significant paralog; % OG Same Sign: n OGs same sign as a percentage of n OGs shared - of the n shared orthogroups, how many have the same sign (DE Up or Down) on their representative logFC in each genome comparison; Gap: The difference between the Jaccard and Weighted Tanimoto score - how much detection agreement is lost once direction/magnitude are also required to agree, where a small gap indicates that the genome pair agrees on the detection and direction/magnitude and a large gap indicates a genome pair agrees on what is significant, but disagrees on the direction/magnitude (or has a different effect size).

| DE Contrast | Genome A | Genome B | Jaccard Score | Tanimoto Score | OGs Shared | % OG Same Sign | Gap |
|---|---|---|---|---|---|---|---|
| SunVsShd_T0 | HapA | HapB | 0.8735 | 0.7702 | 932 | 99.2 | 0.1033 |
|  | HapA | GDDH13 | 0.5923 | 0.5105 | 680 | 97.5 | 0.0818 |
|  | HapB | GDDH13 | 0.6037 | 0.5398 | 690 | 97.4 | 0.0638 |
|  | WA38 | GDDH13 | 0.6118 | 0.545 | 684 | 97.5 | 0.0668 |
|  | HapA | WA38 | 0.8189 | 0.7103 | 886 | 99.2 | 0.1085 |
|  | HapB | WA38 | 0.8168 | 0.6809 | 887 | 98.5 | 0.1359 |
| SunVsShd_2wk | HapA | HapB | 0.8388 | 0.6521 | 359 | 99.2 | 0.1867 |
|  | HapA | GDDH13 | 0.2015 | 0.2438 | 79 | 97.5 | -0.0423 |
|  | HapB | GDDH13 | 0.1931 | 0.2565 | 78 | 98.7 | -0.0635 |
|  | WA38 | GDDH13 | 0.2000 | 0.2113 | 78 | 98.7 | -0.0113 |
|  | HapA | WA38 | 0.7648 | 0.5121 | 335 | 98.5 | 0.2528 |
|  | HapB | WA38 | 0.7384 | 0.4993 | 333 | 98.8 | 0.2391 |
| Sun_2wkvsT0 | HapA | HapB | 0.9187 | 0.8331 | 4982 | 97.8 | 0.0856 |
|  | HapA | GDDH13 | 0.7938 | 0.7012 | 4440 | 95.7 | 0.0926 |
|  | HapB | GDDH13 | 0.7853 | 0.6946 | 4425 | 95.7 | 0.0907 |
|  | WA38 | GDDH13 | 0.7871 | 0.6972 | 4415 | 95.7 | 0.0899 |
|  | HapA | WA38 | 0.887 | 0.7838 | 4874 | 97.2 | 0.1032 |
|  | HapB | WA38 | 0.8837 | 0.7728 | 4877 | 97.3 | 0.1109 |
| Shd_2wkvsT0 | HapA | HapB | 0.911 | 0.8367 | 4563 | 97.8 | 0.0743 |
|  | HapA | GDDH13 | 0.7132 | 0.6855 | 3855 | 94.4 | 0.0278 |
|  | HapB | GDDH13 | 0.7104 | 0.6979 | 3847 | 94.7 | 0.0125 |
|  | WA38 | GDDH13 | 0.7112 | 0.6763 | 3835 | 94.5 | 0.0349 |
|  | HapA | WA38 | 0.8826 | 0.7718 | 4471 | 96.9 | 0.1108 |
|  | HapB | WA38 | 0.8744 | 0.7805 | 4450 | 97.2 | 0.0939 |

We next quantified OGs detected as differentially expressed by other references but absent from each genome’s DE results (**Table S10, File S6**). In SunVsShd_T0, GDDH13 failed to detect 26.2% of the OGs identified by both GS-HapA and ‘WA 38’; GS-HapA and ‘WA 38’ each failed to detect only 3.8-4.4% of those identified by the others. Conversely, approximately 21% of OGs identified as differentially expressed using GDDH13 were not identified using either other reference.

For weighted Tanimoto comparisons, OGs with multiple DEGs (multi-DEGs) were represented by the gene with the largest |logFC| (**Table S11**). In SunVsShd_T0, there were 203, 253, and 240 orthogroups in GDDH13, GS-HapA, and ‘WA 38’, respectively, containing 590, 801, and 754 DEGs. There is no expectation that paralogs in a gene family share a regulatory pattern, so unsurprisingly, 20%-25% of orthogroups with multi-DEGs contained logFC patterns in opposite directions, affecting ∼10% of all genes in multi-DEG orthogroups (**Table S12**). We therefore tested whether representative-selection contributes to the cross-genome disagreement next.

##### 2.7.3.2 Gene Family Functional Profiles - Why Do the Genomes Disagree?

We examined three potential contributors to cross-genome differences in orthogroup profiles: 1) paralog representative selection; 2) reference-generation differences in sequencing technologies and assembly and annotation workflows; and 3) the match between the reference cultivar and the RNA-seq samples.

##### Paralog-Representative Selection Bias

Paralog representative selection contributed to cross-reference disagreement but did not fully explain it. For each pairwise orthogroup comparison, an orthogroup was flagged when significant paralogs within either genome had opposing logFC signs, making the choice of the maximum-|logFC| representative potentially sensitive to paralog selection. For example, in SunVsShd_T0 this affected 20.7–23.7% of multi-paralog orthogroups, with similar incidence across all three genomes (GDDH13: 42/203; GS-HapA: 60/253; ‘WA 38’: 50/240; **Table S12**). The other DE contrasts showed similar patterns but different magnitudes (likely due the different numbers of DE genes identified across different contrasts). We tested whether this discordance is driving the cross-genome disagreement captured by the Jaccard-vs-weighted-Tanimoto Gap (**Table 6**), by recomputing direction agreement (% same sign) and the local Gap separately for orthogroups flagged in at least one of the two genomes being compared versus orthogroups flagged in neither (**Table S13**); note these subset-level values do not sum back to the pair-level Gap reported in **Table 6**, since weighted Tanimoto does not decompose additively.

At the two best-powered contrasts (Sun_2wkvsT0, Shd_2wkvsT0), flagged orthogroups showed a significantly larger local Gap than unflagged orthogroups for all three genome pairs (**Table S13**; bootstrap 95% CI excluding zero in all six cases). In SunVsShd_T0 and SunVsShd_2wk this effect was directionally consistent but did not reach the same level of statistical support, likely reflecting the smaller flagged orthogroup sample sizes at those contrasts (10’s OGs instead of 100’s or 1,000’s). Unflagged orthogroups agreed in logFC sign 98.3–100% of the time across every genome pair and contrast, while flagged orthogroups agreed only 60–88.3% of the time (excepting one contrast/pair cell with only 3 flagged orthogroups). This indicates that representative-selection ambiguity accounts for nearly all logFC sign disagreement between genomes. Residual disagreement among unflagged orthogroups (Gap = 0.050-0.108 across all genome pairs and DE contrasts) indicates that representative-selection ambiguity likely accounts for only part of the observed cross-genome disagreement. To explore additional drivers of cross-genome disagreements, we assessed the reference generation effect.

##### Reference-generation and Cultivar-match Effects

Pairwise disagreement rates were compared to assess potential reference-generation and cultivar-match effects (**Table S14, Table S15**). In SunVsShd_T0, GS-HapA and ‘WA 38’ had similar disagreement rates with GDDH13 (0.388 and 0.408, respectively; p=0.344). Disagreement between GS-HapA and ‘WA 38’, the comparison most directly informative about cultivar matching, was substantially lower (0.181; p < 0.00005) for comparisons with the GDDH13-associated rates (**Table S15**). This pattern was consistent across all four DE contrasts and is compatible with a substantial contribution from reference-generation differences between GDDH13 and the two modern phased references. Residual disagreement between GS-HapA and ‘WA 38’ may additionally reflect cultivar matching, although this contribution cannot be isolated from other reference-specific differences.

## 3. Discussion/Summary

### 3.1 A chromosome-scale, parent-of-origin-resolved ‘Granny Smith’ genome produced through course-based research experience

The parent-of-origin-resolved ‘Granny Smith’ assembly combines chromosome-scale contiguity with high completeness in gene-rich and repetitive regions, as assessed by BUSCO and LAI, respectively. Pedigree-phased SNP markers supported consistent parental assignment to the ‘Cleopatra’-derived HapA and ‘Stone Pippin’-derived HapB. Together with similarly complete annotations and functional information for most predicted genes, these features provide a strong foundation for haplotype-aware analyses.

This resource extends previously published ‘Granny Smith’ genomes. Wang et al ^60^ generated a chromosome-scale haploid/consensus assembly, whereas Watts et al ^61^ reported two haplotype-resolved assemblies. Our pedigree-based evaluation of the latter identified multiple intrachromosomal phase switches, indicating discontinuous parent-of-origin phasing (**Figure 2B**). Reanalysis of their annotations using eudicots_odb10 recovered 95.6% and 95.3% complete BUSCOs, compared with 98.7% and 98.9% for our HapA and HapB. The present resource therefore combines chromosome-scale parental phasing with more complete recovery of conserved genes.

Comparative analyses further distinguished gene-family representation from annotation consistency. Extensive synteny supported conserved chromosome organization, and 9,868 orthogroups were shared among all four ‘Granny Smith’ annotations. However, recovering an orthogroup does not establish that its constituent genes are accurately delineated. CROG analysis complements completeness assessments by comparing annotated gene counts within conserved Rosaceae gene families ^62^. Our ‘Granny Smith’ haplome annotations clustered together near ‘WA 38’ and ‘Honeycrisp’, whereas the Watts et al ^61^ annotations formed a distinct pair separated from most other Malus annotations (**Figure 3A**). Their isolated placement, broad z-score distributions (**Figure S3**), and prominent high-count regions raise substantial concerns about annotation-related distortion of family sizes. Nevertheless, this dendrogram is not a phylogeny, and annotation workflows can drive clustering across taxonomic boundaries ^48^.

The “hot regions” in the Watts et al ^61^ ‘Granny Smith’ annotations therefore warrant investigation before being called gene-family expansions. They represent elevated counts in clustered orthogroups, not contiguous genomic intervals, and could reflect genuine duplications, fragmented or redundant models, retained allelic sequence, or repeat-associated predictions. Protein structure, local synteny, repeat overlap, and transcript or protein support should be examined under comparable annotation criteria before evolutionary inference. Likewise, the 312 orthogroups absent from both ‘Golden Delicious’ T2T annotations require sequence-level examination before being interpreted as gene losses. This caution also applies to our resource: despite close overall haplome agreement, the extreme HapB count in OG2281 and unusually short proteins in several enriched families require targeted review.

Notably, most computational analyses were completed by students during a one-semester American Campus Tree Genomes course. The production of a highly complete, research-grade genome in this setting demonstrates that a structured, mentored course model can contribute valuable community genomic resources while training students through authentic research. The resulting reference therefore advances both haplotype-aware studies of ‘Granny Smith’ and a scalable approach to genomics education.

### 3.2 Reference generation and cultivar matching shape inferred differential expression profiles

A central result of this study is that, although RNA-seq mapping rates differed by only ∼1 percentage point among references, downstream differential-expression and functional-enrichment results differed substantially. The gene families identified as differentially expressed and the biological functions inferred from those genes varied considerably across reference genomes, despite highly similar overall expression trajectories across mapping experiments. This contrast suggests that conventional mapping statistics and broad expression-profile concordance alone may not fully reflect the extent of reference-dependent variation in downstream transcriptomic interpretation.

Functional enrichment revealed a shared core response to sun/heat exposure and postharvest cold storage across all references, but substantial reference-dependent differences emerged beyond this core. Broad responses associated with photosynthesis, chloroplast function, and light exposure were consistently recovered, indicating that major biological signals were relatively robust to reference choice. In contrast, many more specific enriched functions were detected only with a subset of references, and GDDH13 generally recovered the fewest enriched terms. Thus, reference choice had comparatively little influence on the broadest physiological responses but substantially affected the functional resolution of more specific responses.

Orthogroup-level comparisons reinforced this pattern. The two modern phased references, ‘Granny Smith’ and ‘WA 38’, consistently agreed more closely with one another than either did with GDDH13, supporting an important contribution from reference generation, including sequencing technology, assembly quality, phasing, and annotation methodology. Because ‘WA 38’ was generated using approaches closely comparable to those used for ‘Granny Smith’, residual differences between these two references are also compatible with an additional contribution from cultivar matching, although this effect cannot be completely separated from other reference-specific differences.

The influence of reference choice became particularly apparent as the biological contrast became finer. For GS-HapA versus ‘WA 38’, the relative difference between Jaccard and weighted Tanimoto similarity was approximately 11.6-13.3% for the stronger harvest and storage contrasts but increased to 33.1% for the SunVsShd_2wk interaction. At the same time, GDDH13 declined from approximately 4,600 orthogroups containing DE genes in the storage contrasts to only 83 in SunVsShd_2wk, whereas the two modern references declined from roughly 5,000 to ∼385 (**TableS12**). Inspection of individual SunVsShd_2wk orthogroups (**FileS7**) further showed that GS-HapA and ‘WA 38’ often recovered multiple DE genes within the same gene family, whereas GDDH13 recovered only one or none. In many cases, the corresponding GS-HapA and ‘WA 38’ genes also occurred within the same syntenic blocks, supporting their interpretation as likely orthologous counterparts rather than paralogs. Together, these patterns suggest that modern assemblies and annotations likely retain more information as the biological signal becomes weaker or more specific. Reference limitations may therefore become increasingly consequential for interaction effects and other fine-scale regulatory contrasts, where fewer responsive genes contribute to the inferred biological signal.

Orthogroup summarization also influenced estimates of cross-reference concordance. Multi-paralog orthogroups frequently contained significant paralogs with opposing expression responses, making comparisons based on a single representative sensitive to paralog selection. Accounting for this ambiguity reduced some of the apparent disagreement, but did not eliminate the broader differences among references. This distinction is important because high agreement in the direction of shared responses can coexist with substantial differences in which gene-family members are detected and in their estimated effect sizes. Reference-dependent variation therefore reflects both biological and analytical features of the underlying genome representation rather than simple reversals in expression direction.

Together, these findings indicate that improvements in genome assembly and annotation have a clear impact on the resolution of transcriptomic inference, while cultivar matching may provide additional benefits among otherwise comparable modern references. Most importantly, the consequences of reference choice were not uniform across biological questions: broad responses were relatively robust, whereas finer interaction effects were considerably more sensitive to the reference used. This suggests that the value of high-quality, genotype-matched references increases as transcriptomic analyses move from broad physiological responses toward increasingly specific regulatory processes.

### 3.3 Cultivar-matched references recover biologically coherent responses supported by independent metabolomic evidence

The additional functional signals recovered using the cultivar-matched ‘Granny Smith’ references were not simply a larger collection of enriched terms. Several were consistent with metabolomic patterns previously measured in the same experimental system ^35^, providing independent biological support for reference-dependent transcriptomic signals. These included responses associated with photosynthetic pigments, oxidative stress, sterol metabolism, and cuticle biology (**Files S4-S5**), all of which are relevant to the combined effects of preharvest sun exposure and postharvest cold storage. ‘Granny Smith’, particularly GS-HapA, recovered the broadest range of these biologically relevant functions, although ‘WA 38’ also recovered several complementary responses not detected using GDDH13. Thus, the additional resolution provided by modern and cultivar-matched references was associated with pathways independently implicated in the physiology of delayed sun scald rather than simply an increase in the number of enriched functions.

#### 3.3.1 Shared chloroplast and isoprenoid responses

Some of the strongest biological responses were robust across references. At harvest, enrichment of chloroplast and thylakoid functions among genes more highly expressed in sun-exposed peel was consistent with the elevated carotenoids and photosynthetic membrane-associated metabolites previously observed in sun-facing ‘Granny Smith’ peel ^35^. During cold storage, the broad decline in photosystem- and chloroplast-associated expression on both sides of the fruit paralleled the previously reported decline in chlorophyll-related metabolism ^35^. The recovery of these responses across reference genomes indicates that large, coordinated physiological changes can remain readily detectable despite differences in genome representation.

Isoprenoid- and diterpenoid-associated responses were similarly recovered across multiple references and were strongest during storage of previously sun-exposed peel. These pathways have been associated with postharvest chilling responses in apple and are consistent with the substantial metabolic remodeling that occurs as sun-exposed fruit enters cold storage ^35^. Their broad cross-reference recovery further distinguishes these robust responses from the more reference-sensitive functional signals described below.

#### 3.3.2 Reference-dependent oxidative-stress, sterol, and cuticle responses

Other biologically relevant responses depended more strongly on the reference used. Oxidative-stress enrichment at harvest was detected with GS-HapA but not with GDDH13 or ‘WA 38’. This signal is consistent with the elevated E-2-pentenal and E-2-hexenal previously measured in sun-facing peel ^35^, compounds associated with lipid peroxidation and oxidative stress ^63,64^. ‘WA 38’ provided complementary evidence through enrichment of the ascorbate-glutathione cycle during storage, implicated in sun-stress mediation ^65^, further supporting oxidative stress as an important component of the response to prior sun exposure. The reference-dependent recovery of these related pathways illustrates how different genome representations can alter the level of functional detail obtained from the same transcriptomic dataset.

Sterol metabolism showed a similar pattern. Sterol-associated enrichment was recovered with both ‘Granny Smith’ and ‘WA 38’ at harvest, but the sun-exposure × storage interaction was detected only with the ‘Granny Smith’ references. The positive interaction indicates a more positive storage-associated expression change in sun-exposed than shade-exposed peel, rather than necessarily a reversal of the harvest-time difference. Alongside the metabolite patterns reported by McTavish et al ^35^, this finding supports a role for sterol-related responses during storage that depend on prior sun exposure.

Cuticle-associated functions provided another example of additional biological resolution with the matched reference. Lipid-transfer and cutin-biosynthetic processes became prominent during storage and were recovered predominantly, or in some cases exclusively, using the ‘Granny Smith’ haplomes. These transcriptomic signals are consistent with the elevated hydroxycinnamoyl-ester cuticle components previously observed in sun-exposed peel ^35^. Together, the oxidative-stress, sterol, and cuticle responses provide independent examples in which reference-dependent functional enrichment corresponds to metabolomic features already associated with sun exposure and postharvest physiology.

#### 3.3.3 Haplotype-dependent functional enrichment

Functional inference also differed between the two ‘Granny Smith’ haplomes, with each recovering biological processes not detected using the other. These differences demonstrate that downstream interpretation can be haplome-sensitive even within the same cultivar and reinforce that the broader enrichment recovered using GS-HapA did not encompass every response detected with GS-HapB or the non-native references. Such differences may reflect sequence divergence, gene-content and annotation differences, copy-number variation, mapping behavior, or genuine haplotype-specific regulation; the ∼3,600 additional annotated genes in HapB may also contribute.

Because reads were mapped independently to each haplome, however, these patterns should not be interpreted as direct evidence of allele-specific expression. Rather, they show that the choice of haplome itself can influence the functional profile inferred from the same RNA-seq dataset. Direct allele- and paralog-specific expression analyses will be required to distinguish genuine regulatory differences between the two ‘Granny Smith’ haplotypes from effects introduced by genome representation and annotation.

### 3.4 Reference choice shapes transcriptomic interpretation of delayed sun scald in ‘Granny Smith’

Reference choice influenced transcriptomic interpretation far beyond the modest differences observed in read mapping. Modern phased references showed greater agreement in gene-family responses than GDDH13, while the cultivar-matched ‘Granny Smith’ references recovered additional functional signals, particularly for the subtle sun-exposure × storage interaction. These results indicate that both reference generation and genotype matching can affect the biological resolution of transcriptomic analyses.

For delayed sun scald, several signals recovered with the matched reference, including oxidative-stress, sterol, and cuticle-associated responses, were consistent with metabolomic patterns observed in the same experimental system. More broadly, reference selection should therefore be treated as part of experimental design, especially in highly heterozygous crops where subtle, genotype-specific responses are the biological target.

## 4. Materials and Methods

Markdown documentation of the full methods, scripts (including all used parameters), and small intermediate result files are available from the project GitLab repository (https://gitlab.com/ficklinlab-public/gs-genome-project). The ‘Granny Smith’ genome was developed as part of the American Campus Tree Genomes (ACTG) project, with most computational steps carried out by students during a one-semester course.

### 4.1 Sample collection, DNA isolation, and sequencing

Young leaves were collected from one greenhouse-grown *Malus domestica* ‘Granny Smith’ on M-7 rootstock, flash-frozen, and stored at -80 °C until DNA isolation. DNA extraction, sequencing library preparation, and sequencing was performed following the same protocols used to generate the ‘d’Anjou’ pear ^66^ and ‘WA 38’ apple ^48^ genomes. Briefly, high-molecular-weight DNA supported PacBio HiFi sequencing, an Omni-C library provided long-range information for scaffolding, and CTAB-extracted DNA ^67^ was used for Illumina PCR-free shotgun sequencing to estimate genome size and heterozygosity.

### 4.2 Sequence quality assessment and genome complexity analysis

PacBio HiFi, Omni-C, and Illumina shotgun reads were assessed with FastQC ^68^, including shotgun reads before and after trimming. Jellyfish ^69^ k-mer profiles from HiFi and untrimmed shotgun reads were interpreted with GenomeScope 2.0 ^70^ to estimate genome size, heterozygosity, and repeat content.

### 4.3 Genome Assembly

#### Contig assemblies

PacBio HiFi and Omni-C reads were assembled with Hifiasm ^71^ into phased Hap1 and Hap2 contigs. These outputs are termed the contig assemblies and provided the sequence templates for scaffolding.

#### Scaffold assemblies

Omni-C reads were mapped separately to each contig assembly, checked for coverage and mapping quality, and used by YaHS ^72^ to order and orient contigs. Hi-C contact maps generated with Juicer were inspected in Juicebox ^73,74^ for misassemblies; none were detected. The reviewed YaHS outputs are termed the scaffold assemblies and were used for all subsequent assembly processing.

#### Final assemblies and haplotype phasing

Scaffolds were aligned to the ‘WA 38’ HapA genome ^48^ to assign chromosome names and orientations, then unplaced scaffolds were screened to remove plastid and microbial contamination. Because Hifiasm labels do not contain parental origin across chromosomes, array-marker and pedigree information was used to assign chromosome-scale haplotypes ^75^. Chromosomes inherited from ‘Cleopatra’ were grouped as HapA and those from ‘Stone Pippin’ as HapB; unplaced scaffolds retained their original Hifiasm assignment. These cleaned, parentally phased HapA and HapB sequences are termed the final assemblies and were used for annotation.

#### Assembly assessment

Assemblathon statistics ^76^ and BUSCO completeness ^77^ (against eudicots_odb10) were evaluated for the contig, scaffold, and final assemblies. Nucmer/dnadiff whole-genome alignments of both initial and final haplomes quantified sequence similarity and structural variation ^78^.

### 4.4 Structural and Functional Annotation

#### Repeats and telomeres

EDTA ^79^, aided by ‘WA 38’ coding sequences, generated a transposable-element library for each haplome; RepeatMasker ^80^ extended repeat detection and produced masked assemblies. Telomeric repeats were identified with tidk ^81^.

#### BRAKER2 and BRAKER3

Each haplome was annotated independently with BRAKER2 ^82^ and BRAKER3 ^83^ using the same RNA-seq and protein evidence previously employed for annotation of the ‘WA 38’ genome ^48^. BRAKER2 retained the established apple workflow, in which evidence-specific predictions are combined with TSEBRA ^84^, providing students with a clearer view of how different evidence types influence gene-model prediction. BRAKER3 is included as a newer integrated strategy and has also produced high-quality annotations for another ACTG genome, *Carya glabra* ^85^. BUSCO, chromosome-wide IGV inspection ^86,87^, sequence/domain searches, and PlantTribes2 ^62^ orthogroup comparisons showed that BRAKER3 models were generally more complete (**FileS8**-Example1-3) and better supported, so BRAKER3 was selected as the baseline.

#### Integrated BRAKER2 + BRAKER3 annotation

BRAKER3-only models lacking both InterPro ^88^ and orthogroup support were removed, whereas BRAKER2-only models supported by both (**FileS8**-Example4) were added. Manual review addressed a recurrent BRAKER3 error in which adjacent tandem duplicates were fused (**FileS8**-Example5): when BRAKER2 and RNA-seq coverage supported separate genes, the fused model was replaced by the corresponding BRAKER2 models.

#### PASA refinement and curation

*De novo* and haplome-guided transcriptome assemblies were supplied with the reconciled models to PASA for transcript-supported refinement and UTR annotation ^89^. Three update rounds were completed because improvement had plateaued by the third round. IGV review corrected PASA changes unsupported by RNA-seq, especially erroneous merging of adjacent genes. Models largely overlapping repeats were removed, and models containing non-overlapping transcript isoforms were split. Final models were named according to AgBioData recommendations ^90^ and assessed with AGAT ^91^ and BUSCO.

#### Functional annotation

The final HapA and HapB gene sets were functionally annotated with AnnoTater (https://github.com/SystemsGenetics/AnnoTater) using InterProScan and similarity searches against RefSeq and UniProtKB/Swiss-Prot.

### 4.5 Comparative analysis

GENESPACE ^92^ analyses evaluated chromosome-scale conservation among ‘Granny Smith’, high-quality domesticated apple genomes, representative wild *Malus* species, and *Pyrus communis* ‘d’Anjou’ as an outgroup. PlantTribes2 ^62^ assigned HapA and HapB proteins to 26Gv2.0 orthogroups. These assignments were incorporated into the ‘WA 38’ pome-fruit dataset to run a CROG analysis to identify shared and cultivar-specific orthogroups and to compare normalized core orthogroup gene counts across 27 genomes, revealing variation in gene-family representation among genomes ^48^.

### 4.6 Organelle Genome Assembly and Annotation

Following Zhang et al ^48^, adapter-trimmed Illumina reads were assembled with NOVOPlasty ^93^ to recover the chloroplast genome, whereas Hifiasm contigs were assembled with MitoHiFi ^94^ to recover the mitochondrial genome. Both organelle genomes were annotated with GeSeq ^95^.

### 4.7 Comparative RNA-Seq Mapping Experiment

#### 4.7.1 Experimental design, tissue collection, RNA extraction, quality control, and transcriptome sequencing

‘Granny Smith’ fruit and peel tissue were used from a previously published study ^35^ on field irradiance/heat stress and postharvest chilling stress (full experimental details therein). Fruits were harvested at commercial maturity from a commercial orchard near Mattawa, WA, USA, selecting fruit with clear sun-exposed/shaded contrast. Pooled peel samples (3 biological replicates of 6 apples each) were collected from the sun-exposed and shaded sides at 0, 14, 30, 60, and 180 d postharvest, flash-frozen, ground, and stored at −80°C until RNA extraction using a modified CTAB protocol ^96^. RNA quantity and purity were assessed following Honaas et al ^43^; only RNA meeting A260/A280 ≈ 2.0 and RIN ≥ 8.0 was used. Libraries were prepared at the Penn State Genomics Core Facility (University Park, PA) using Illumina’s TruSeq Stranded mRNA Library Prep kit and sequenced 150 bp single-end on an Illumina HiSeq 2500 (Rapid Mode).

#### 4.7.2 Transcriptome data quality control, reference selection, and read mapping

Raw reads were processed with GEMmaker ^97^ using STAR ^98^. To compare read mapping to matched versus non-matched cultivar references, reads were independently mapped to four references 1) GDDH13 v1.1 ^42^, 2) ‘WA 38’ HapA ^48^, 3) ‘Granny Smith’ (GS) HapA, and 4) GS HapB. Mapping statistics were generated with MultiQC (v1.11, **File S2**).

#### 4.7.3 Differential Expression and Functional Enrichment

Differential Expression (DE) analysis focused on expression changes from harvest (T0) to 14d (days; 2wk) postharvest storage on each fruit side, and the fruit-side × time interaction, based on prior evidence that the most metabolically relevant sun scald-related changes occur in this window ^35^. For each reference genome, DE was evaluated in edgeR ^99^ (v4.8.1), with fruit side as a factor and time as a covariate. Dispersion was modeled with ‘glmQLFit()’ ^100,101^ (a conservative choice for n=3/group), and contrasts tested with ‘glmQLTest()’. Significant DE was thresholded at FDR < 0.05 and |log2FC| > 1.

DE genes were hierarchically clustered by expression profile (correlation-based distance, Ward’s D2 linkage). Since unsupervised clustering does not align clusters across experiments, per-cluster centroid profiles (mean z-scaled expression per sample) were Pearson-correlated between a reference experiment (GS-HapA) and each other experiment, and clusters matched via the Hungarian algorithm ^102,103^ (R package “clue,” ‘solve_LSAP()’) for globally optimal one-to-one assignment; the “padded” option accommodated differing per-experiment cluster counts (§3.0). GO enrichment was performed per cluster with FUNC-E (v2.0.1) ^104^.

#### 4.7.4 Cross Genome Comparisons of Differential Expression and Functional Profile

DE results and functional profiles were compared across GDDH13, GS-HapA, and ‘WA 38’ using PlantTribes2 ^62^ 26Gv2.0 orthogroup assignments of the longest protein isoform, with agreement quantified by the Jaccard index (detection agreement) and a magnitude/direction-weighted Tanimoto score. Signal reproducibility was tested as whether a significant orthogroup in one genome was also independently significant in both others. Genes unassigned to orthogroups were excluded (**Table S16**).

Where an orthogroup contained multiple significant DE paralogs, paralogs were collapsed to the single paralog with maximum absolute logFC (not the mean, since averaging opposite-signed paralogs can produce a sub-threshold value even when each individually exceeds |logFC| ≥ 1; **Table S11**). Because this representative spans both directions, an orthogroup can be counted in a genome’s “Up” set while its representative is a larger magnitude Down paralog, or vice versa; every instance was flagged.

#### 4.7.5 Investigation of why differences across genomes exist

We investigated two non-exclusive sources of the observed cross-genome disagreement: 1) a methodological artifact of the paralog-representative selection, and 2) genuine differences between genomes arising from reference assembly/annotation quality and/or cultivar match to the RNA-seq source.

##### 4.7.5.1 Methodological - Paralog representative selection bias

Anchoring the weighted Tanimoto score to the single strongest signal in a multi-paralog orthogroup is arbitrary when paralogs are independently significant in opposite directions (**Table S12**). An orthogroup was flagged as discordant for a given pairwise comparison if either genome had opposite-signed paralogs for that orthogroup (**Table S9**); and direction agreement and a local Jaccard/Tanimoto gap were then recomputed separately within the flagged and unflagged subsets of each genome pair (**Table S13**). These subset-level values do not sum to the pair-level Gap in **Table 6**, since weighted Tanimoto does not decompose additively. This methodological effect explained part, but not all, of the observed disagreement, motivating the investigation below.

##### 4.7.5.2 Reference genome quality and native cultivar match

We investigated whether the remaining disagreement reflects a pipeline/assembly-quality effect, independent of cultivar match, versus a native-match/read-mapping-specificity effect, by comparing pairwise disagreement rates across the three genome pairs: GDDH13 × ‘WA 38’ and GDDH13 × GS-HapA both change assembly pipeline and phasing; GS-HapA × ‘WA 38’ isolates native cultivar match alone, holding pipeline and phasing constant. As GDDH13 contributes to two of the three comparisons, orthogroup-level disagreement outcomes are correlated across comparisons rather than independent, we used an orthogroup-level permutation test. For each pair of comparisons being contrasted, orthogroups present in both had their disagreement outcome randomly reassigned between them (20,000 permutations), generating an empirical null distribution for the observed rate difference (two-sided p-value), applied to all three pairwise comparisons at each of the four contrasts.

## 5. Acknowledgements

The authors would like to acknowledge the American Campus Tree Genomes Project for designing course materials and running the course, Christine McTavish for providing images and project information details, and Cameron Peace and Nick Howard for their roles in generating the phased SNP array data and ‘Granny Smith’ parent identification. ClaudeAI Sonnet 5.5 was used to generate Alt Text captions for figures, captions were reviewed by H.H. and H.Z.

## 6. Contributions

H.H., H.Z., Z.A., A.A., A.D., E-O.I., H.K., I.L., D.L., C.M., B.M., T.M., B.N., O.O., S.P., & E.S. performed the genome assembly, initial annotation, and drafting of the manuscript. H.Z., I.K., and S.F. instructed the ACTG course. H.H., H.Z., I.L., H.K., D.L., and A.A. performed manual curation of annotation. H.Z. finalized the structural and functional annotation and performed comparative analyses. H.H. performed the mapping experiment, case study analysis, and wrote the final manuscript draft. S.V. provided curated parental phased SNPs for parental phasing confirmation. D.R. designed the postharvest storage experiment and provided tissue samples for RNA-Seq. A.H. and L.H. provided funding. All authors reviewed and edited the final manuscript.

## 7. Data Availability Statement

Raw reads generated for this project are publicly available at NCBI under BioProject: PRJNA1562996. Genome assembly and annotation are available on GDR: https://www.rosaceae.org. Markdown documentation of the full methods, scripts (including all used parameters), and small intermediate result files are available from the project GitLab repository (https://gitlab.com/ficklinlab-public/gs-genome-project).

